# Complete microbial genomes for public health in Australia and Southwest Pacific

**DOI:** 10.1101/829663

**Authors:** Sarah L. Baines, Anders Gonçalves da Silva, Glen Carter, Amy V. Jennison, Irani Rathnayake, Rikki M. Graham, Vitali Sintchenko, Qinning Wang, Rebecca J. Rockett, Verlaine J. Timms, Elena Martinez, Susan Ballard, Takehiro Tomita, Nicole Isles, Kristy A. Horan, William Pitchers, Timothy P. Stinear, Deborah A. Williamson, Benjamin P. Howden, Communicable Diseases Genomics Network (CDGN), Torsten Seemann

## Abstract

Complete genomes of microbial pathogens are essential for the phylogenomic analyses that increasingly underpin core public health lab activities. Here, we present complete genomes of pathogen strains of regional importance to the Southwest Pacific and Australia. These enrich the catalogue of globally available complete genomes for public health while providing valuable strains to regional public health labs.

**Announcement:** Whole-genome sequence (WGS) data is increasingly important in public health microbiology (1–4). The data can be used to replicate many of the basic bacterial sub-typing approaches, as well as support epidemiological investigations, such as surveillance and outbreak investigation (5–7). The appeal of WGS data comes from the promise of a single workflow to process all microbial pathogens that can provide easily portable data that promotes deeper integration of surveillance and investigation efforts across jurisdictions. This promise is leading to a concerted effort to move microbial public health to a primarily genome-based workflow at numerous jurisdictions (8–10), including Australia (11).

---

Essential to the success of this transition to a genomics workflow is the need to develop catalogues of high-quality complete genomes of microbial pathogens (12). Complete genomes of bacterial pathogens can provide valuable insight, for instance, into the genomic context of virulence genes and antimicrobial resistance genes (13), and their possible mechanisms of actions. More importantly, complete genomes are essential for generating accurate phylogenomic analyses, a core requirement for public health surveillance and outbreak responses. In this setting, complete genomes provide valuable context to identify variable genomic regions across samples in a given study in a computationally efficient manner (14–16).

However, pathogenic bacteria are not generally composed of uniform panmictic populations. Instead, they represent numerous diverse clades, with many being endemic to particular regions or jurisdictions (17–23). This inherent pathogen population structure poses a challenge to a successful transition to genomics in public health labs because it can significantly reduce the resolution of phylogenomic analyses by affecting the identification of genetic variants (24). Thus, catalogues of complete genomes will only be effective in supporting a transition to genomics in public health labs if they are rich in endemic strains. Here, we present 26 complete genomes of microbial pathogens of regional significance to the Southwest Pacific and Australia (Table 1). We will continue to build on this resource as further strains are sequenced and assembled.

**Table 1:**
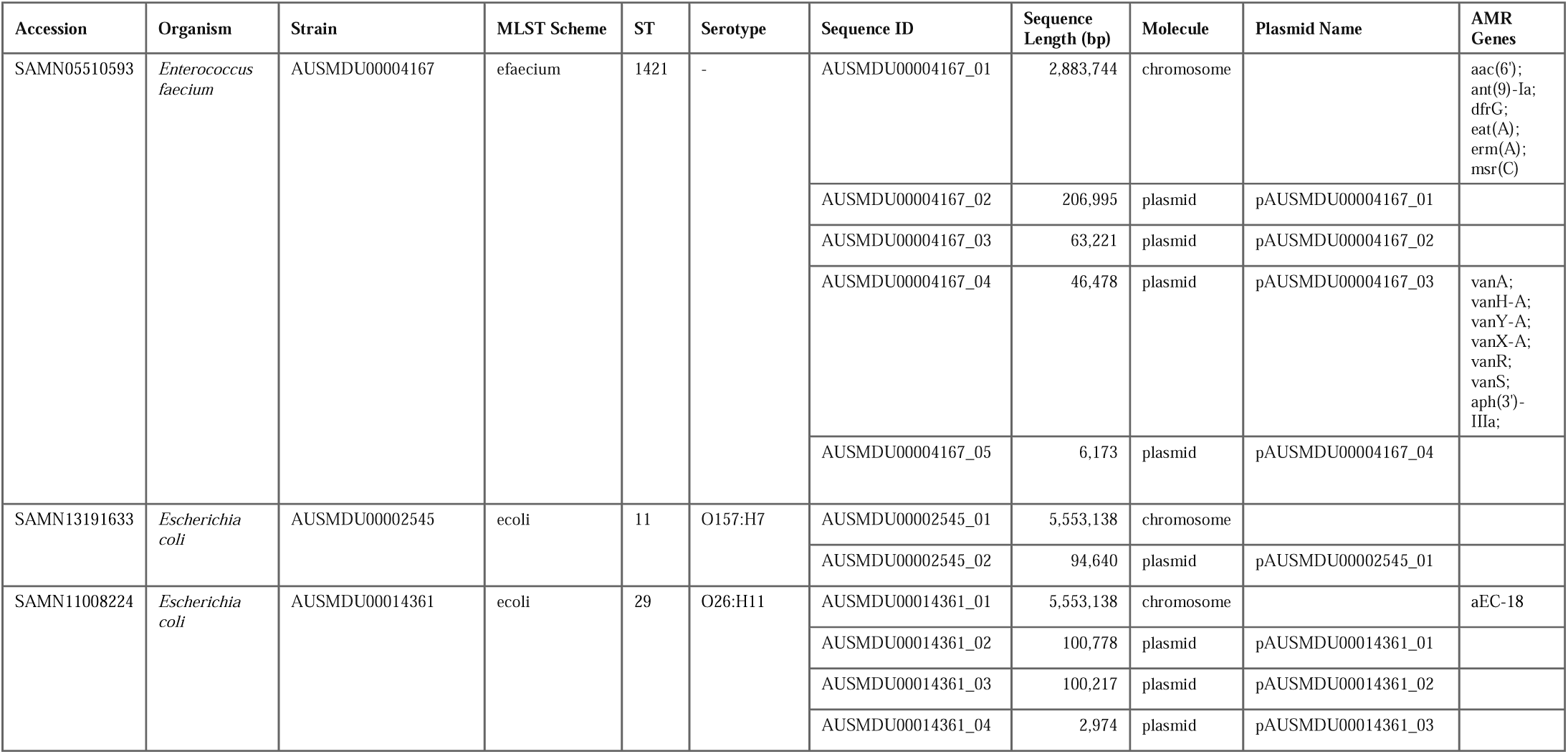
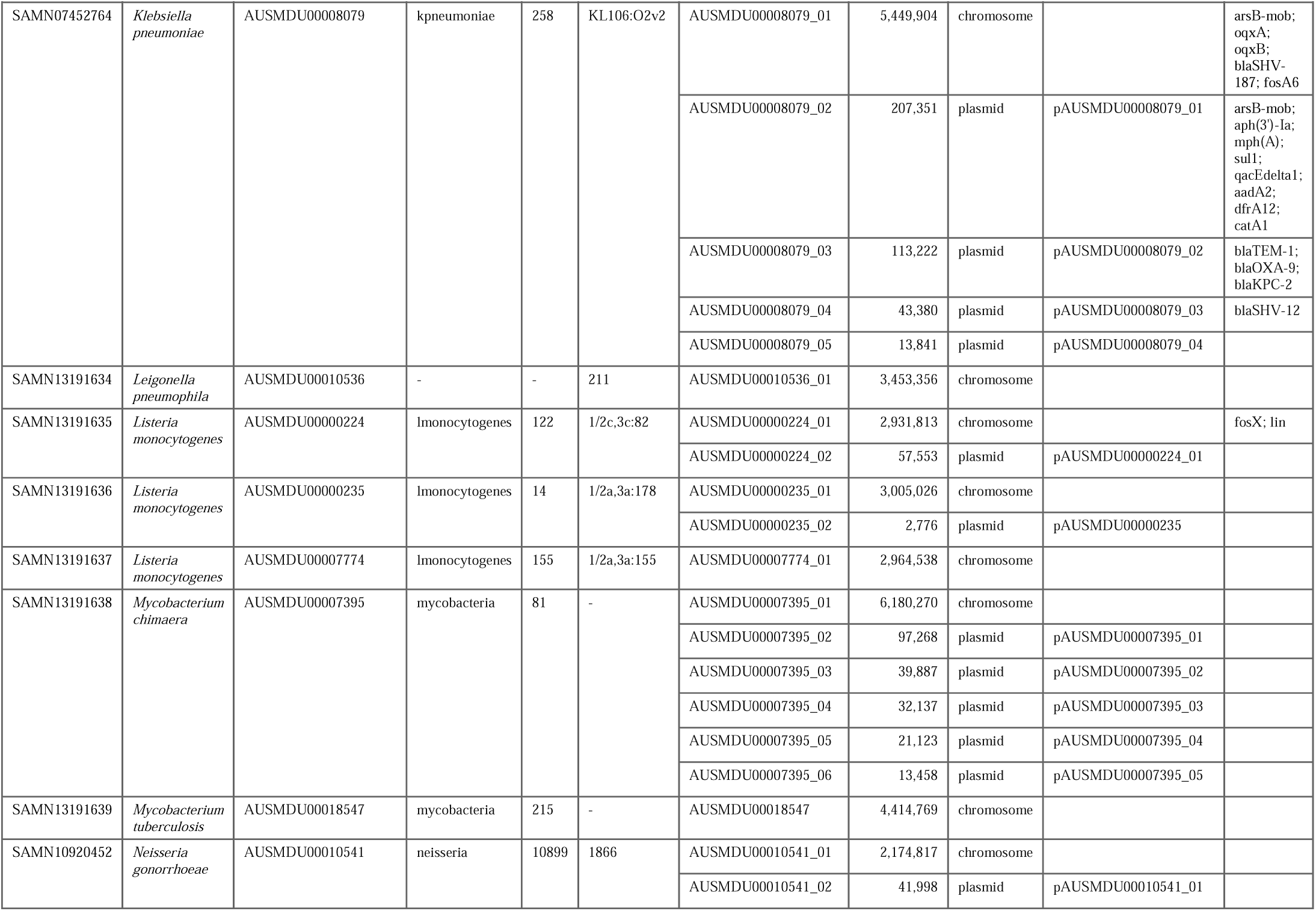
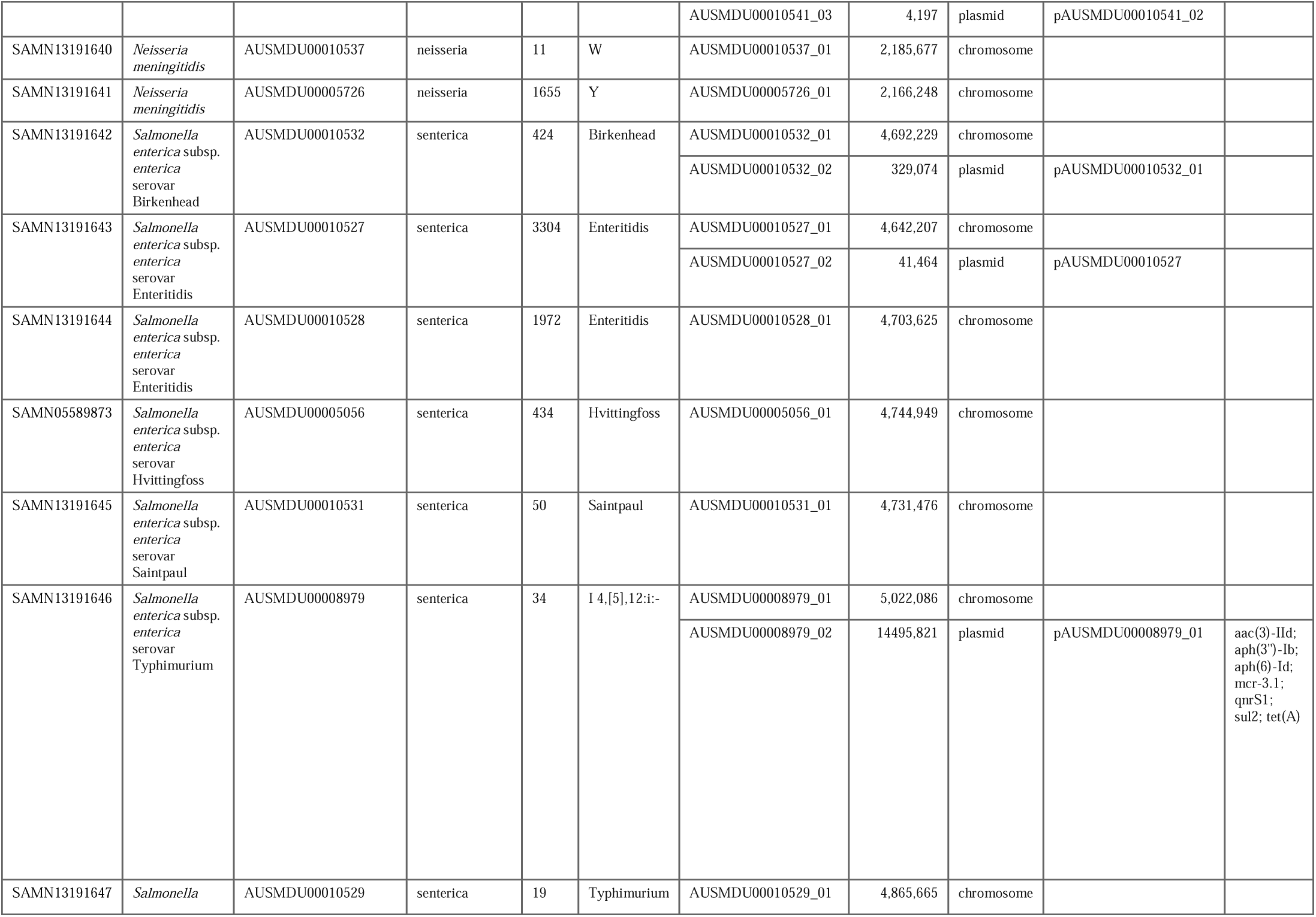
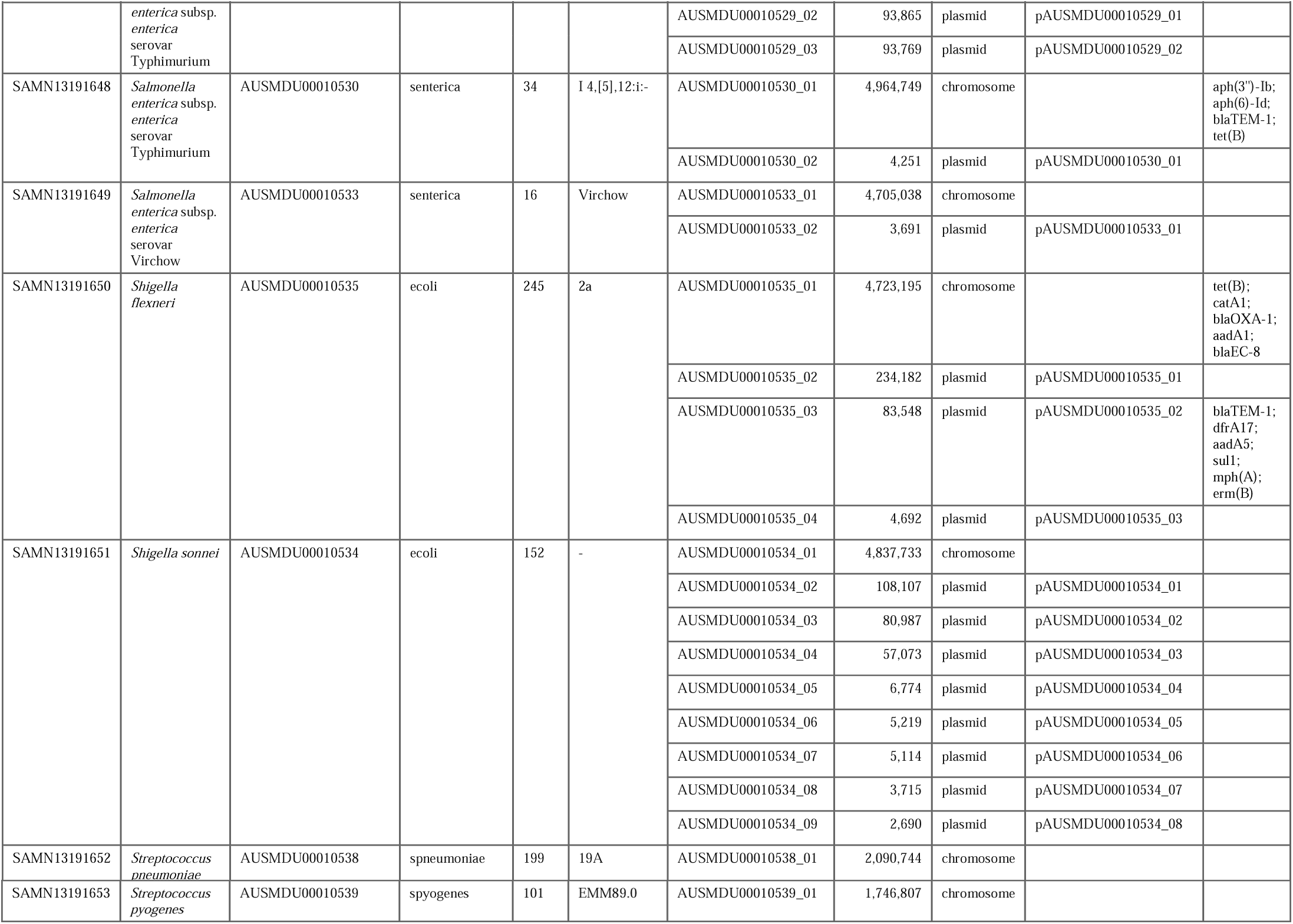
Sumamry data of complete genomes.

Strains were selected to represent a broad range of organisms of importance for public health in Australia. These included foodborne (e.g., *Listeria monocytogenes*, and *Salmonella enterica*), waterborne (e.g., *Legionella pneumophila*), sexually transmitted (e.g., *Neisseria gonorrhoeae*), and other pathogens of public health importance (e.g., *Klebsiella pneumoniae, Mycobacterium* sp., *N. meningitidis*). In some cases, we chose the strains because of their relevance to local surveillance and outbreak requirements as well as their virulence or antimicrobial resistance (AMR) phenotypes (e.g., Colistin-resistant *Salmonella enterica* and Vancomycin-resistant *Enterococcus faecium*). We deposited pure cultures of all strains in the Microbiological Diagnostic Unit Public Health Laboratory (MDU PHL) Reference Culture Collection.

All strains were grown in appropriate media for the organism following standard laboratory protocols. Whole genomic DNA was extracted using various methods, selected based on the particular species to ensure high-quality DNA for long-read sequencing. Samples were then sequenced on either the Illumina NextSeq500/550 or MiSeq platforms to generate short-read sequence data, and either the PacBio RS-II or Oxford Nanopore GridIon platforms to generate long-read sequence data. DNA extraction, library preparation and sequencing workflow are illustrated elsewhere (25).

Before *de novo* assembly, all sequence data were quality controlled to ensure sufficient depth (>40X), basecall quality (mean >Q30), and that the sequence data represented the expected organism based on a kraken2 (26) analysis. Confirmation of the *Shigella sonnei* identification was done by phylogenetic analysis of the isolate against samples described in (27). All genomes were assembled using four different approaches, with the resulting assemblies compared for structural consistency and recovery of small replicons (e.g., plasmids). When consistent, we prioritised assembly approaches in the following order: Hybrid assembly with unicycler (28) v0.4.6 and v0.4.8b, long-read only assembly with HGAP3 (29) (SMRTPortal v2.3.0), long-read only assembly with Canu v1.8 (30), and long-read only assembly with unicycler v0.4.6 and v0.4.8b. The selected assembly was further error corrected as outlined elsewhere. Strain specific assembly details can be found in (25).

*In silico* multilocus sequence typing (MLST) was performed using mlst (31) v2.11 with the pubMLST (32) schemes. AMR genes were detected using abricate (33) v0.8.13 with the NCBI database (34). AMR SNPs were detected using Ariba (35) v2.14.4 using the CARD database (36) and the built-in TB database. *In silico* serotyping was performed using emmtyper (37) v0.1.0 for *Streptococcus pyogenes*, kleborate (38) v0.3.0 for *Klebsiella pneumoniae*, legsta (39) v0.3.2 for *Legionella pneumophilia*, lissero (40) v0.2 for *Listeria monocytogenes*, meningotype (41) v0.8.2-beta for *N. meningitidis*, ngmaster (42) v0.5.5 for *N. gonorrhoeae*, seroba (43) v1.0.1 for *Streptococcus pneumoniae*, shigatyper (44) v1.0.5 for *Shigella spp*., sistr (45) v1.0.2 for *Salmonella enterica*, and srst2 (46) v0.2.0 using the EcOH DB for *Escherichia coli*.

## Data availability

All sequencing data and assemblies have been deposited in NCBI under the BioProject no. PRJNA556438.

## Acknowledgments

This study was funded by a grant from the Department of Health of the Australian Federal Government to the Communicable Diseases Genomics Network (CDGN). The authors and the CDGN would like to thank and acknowledge the pathology and state reference laboratory staff that performed the original isolation and characterisation of the strains included in this work.

